# microRNA-mRNA interaction identification in Wilms tumor using principal component analysis based unsupervised feature extraction

**DOI:** 10.1101/059295

**Authors:** Y-h. Taguchi

**Affiliations:** Department of Physics, Chuo University, Tokyo 112–8551, Japan.

**Keywords:** principal component analysis, feature selection, Wilms tumor

## Abstract

Wilms tumor is one of lethal child renal cancers, for which no known disease causing mechanisms exist. In this paper, we tried to identify possible disease causing microRNA(miRNA)-mRNA pairs (interactions) by analyzing (partially matched) miRNA/mRNA gene expression profiles with the recently proposed principal component analysis based unsupervised feature extraction. It successfully identified multiple miRNA-mRNA pairs whose biological natures are convincing. Correlation coefficients between miRNA and mRNA expression in matched parts of profiles turned out to be significantly negative. Constructed miRNA-mRNA network will be a key to understand Wilms tumor causing mechanisms.

## I. INTRODUCTION

Wilms tumor [1] is one of lethal child renal tumor whose disease causing mechanism is unknown. Especially, bilateral Wilms tumor is difficult to treat [2]. In order to develop the effective therapy, it is critically important to understand how Wilms tumor develops from normal kidney. In this regards, potential role of microRNA(miRNA) in Wilms tumor development recently collected broad interests [3], [4], [5], [6], [7].

Among those researches, although Luding et al [3] successfully identified feasible miRNA-mRNA pairs, from the methodological point views, there remains some possibilities to be improved. For example, they employed non-adjusted *P*-values to identify differently expressed miRNA/mRNAs between Wilms tumor and healthy control. Since the number of miRNAs/mRNAs considered were huge, it is better to employ adjusted *P*-values to identify significant changes. They also used fold changes (FC) to screen mRNA/miRNAs. They employed FC> 2 as threshold. Since it is a very standard criterion, none would be oppose to this selection, but why *two* is reasonable number? Are there any biological reasoning to do so? Finally, in their Table 1, they listed top most up-/down-regulated 15 miRNAs. Although they also looked so reasonable from the biological point of views, why did they select specifically not 10 or 20, but 15 miRNAs? Possibly, the answers are simple; these criteria were employed since the outcomes are biologically feasible. Since none of criteria employed are unrealistic, if the outcome is feasible, there are no needs to criticize it. However, if we could get similar outcomes without optimizing various criteria, it is more hopeful.

In the previous study [8], the recently proposed principal component analysis (PCA) based unsupervised feature extraction (FE) was successfully applied to miRNA-mRNA interaction identification in various cancers. It worked pretty well although it was not modified from samples to samples (from cohorts to cohorts) so as to get feasible results, but employed single common criterion to identify feasible miRNA-mRNA pairs. In this paper, the almost same strategy was applied to data set used by Luding et al [3], and it turned out to work well for these data sets; for example, top 15 down-regulated miRNAs in Wilms tumor compared with normal kidney shown in thier Table 1 were almost identified without optimizing almost anything (no FC threshold, no specified number of top ranked miRNAs, and adjusted *P*-values are used).

## II. METHODS

Overall study work flow is shown in Fig. 1.

**Figure 1.**
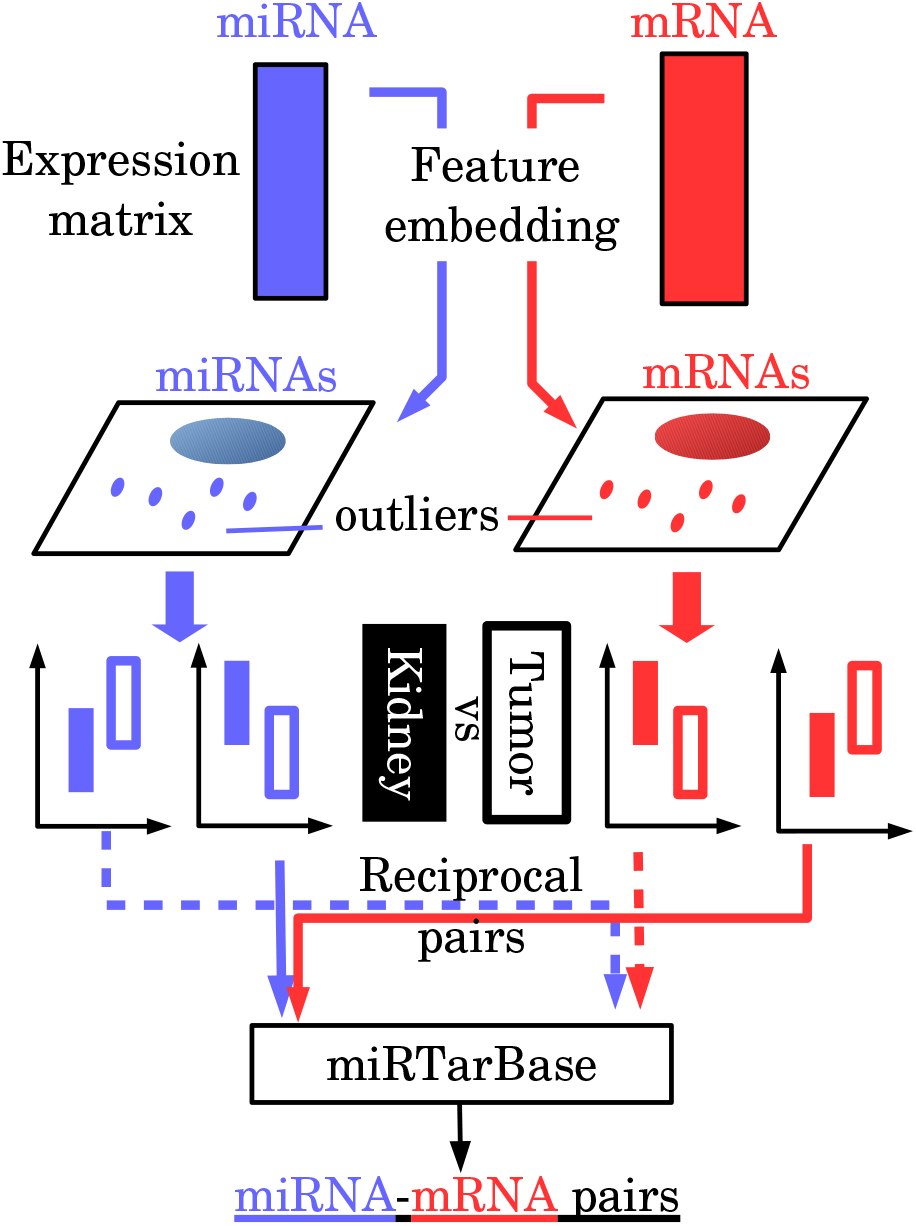
Workflow of this study.miRNA/mRNA expression profiles were separately embedded into low dimensional space by PCA (feature embedding). After identifying PCs used for FE, outlier miRNAs/mRNAs are selected. miRNAs/mRNAs exhibiting significant differential expression between tumor and normal kidney were further selected among those selected as outliers, and pairs associated with reciprocal expression were compared with those listed in miRTarBase.

### A. mRNA/miRNA expression profiles

mRNA/miRNA expression profiles were extracted from GEO using GEO ID GSE66405/GSE57370, respectively (processed data within sampletable). getGEO function implemented in GEOquery [9] package (Biocounductor) were used to load profiles into R [10]. Each profile is normalized so as to have zero mean and unit variance within each sample. mRNA samples composed of 28 Wilms tumor samples and 4 normal kidney samples. miRNA samples composed of 62 Wilms tumor samples and 4 normal kidney samples. No subtype information was used in this study. Thirty two samples having mRNA expression profiles also have miRNA expression (matched samples).

### B. *PCA based unsupervised FE*

Although PCA based unsupervised FE was successfully applied to various bioinformatics problems [11], [12], [13], [14], [15], [16], [17], [18], [19], [20], [21], [22], [23], [24], [25], [26], we briefly describe the outline of this methodology. Let *xij* be the expression of the ith mRNA/miRNA of the *j*th sample. The elements *x_ij_* are contained in a matrix *X*, and we standardize the columns in the matrix. In contrast to standard PCA, which embeds the samples, PCA based unsupervised FE embeds the genes (miRNAs or mRNAs).

Then *k*th principal component (PC) score *u_ki_* attributed to the *i*th gene is computed as an element of the eigenvector u*_k_* of the gram matrix *G* ≡ *XX^T^*,

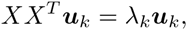

where the eigenvalues λ*_k_* are ordered such that λ_*k*+1_ < λ*_k_*.

Because we have

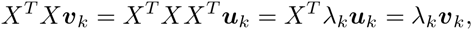

the *k*th PC loading *v_k_j* attributed to the jth sample is computed as an element of v*_k_* = *X^T^* u*_k_*, which is an eigenvector of the matrix *X^T^X*. After identifying a set Ω*_k_* of PCs with distinctly different loadings between tumors and normal tissues (*t* test, *P* < 0.05), the outlier genes are identified by a *χ* squared distribution, assuming a Gaussian distribution of the PC scores:

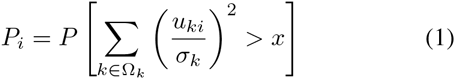

where *P*[> *x*] is the cumulative probability of the *χ* squared distribution, where the argument exceeds *x* and *σ_k_* is the standard deviation of the kth PC scores. Then, if the Benjamini-Hochberg (BH) criterion [27]-adjusted *P_i_* is below 0.01, gene *i* is identified as an outlier.

### C. Identification of significant miRNA-mRNA pairs

Some of the mRNA/miRNAs selected as outliers by the PCA-based unsupervised FE showed significant up-/down-regulation between normal control tissues and tumors (BH criterion [27]-adjusted P < 0.05, *t* test). The list of conserved target genes of each miRNA was then obtained from mirTarBase [28], and the miRNA–mRNA pairs associated with reciprocal regulation and identified by miRTarBase were selected.

### D. Discrimination between Wilms tumor and normal kidney

Discrimination was performed by linear discriminant analysis (LDA) using PCA [23], [22], [21]; The LDA was performed by the lda function in R [10]. In this analysis, the PC loadings were recomputed using only the mRNAs or miRNAs selected by the PCA-based unsupervised FE. The recomputed loadings were then attributed to samples. The leave-one-out cross validation was employed since we set CV=T. We also weighted both classes equally by setting prior=rep(1/2,2). The first *L* PC loadings were used for discrimination, and the optimal *L* for each cancer was found by trial-and-error.

## III. RESULTS

### A. PCA based unsupervised FE applied to miRNA/miRNA expression profile

miRNA and mRNA expression profiles were separately embedded into low dimensional space by PCA (feature embedding). Then, we found that the first to the third PC loadings for mRNA and the second as well as the forth to the seventh PC loadings for miRNA were significantly distinct between Wilms tumor and normal kidney, respectively (*t* test, *P*-values < 0.05, see Methods). Using these PCs, we computed adjusted *P*-values assuming *χ* square distributions for PC scores attributed to each mRNA/miRNAs. After identifying outliers (adjusted *P*-values < 0.01, see Methods), we have gotten 55 miRNAs and 1114 probes attributed to each mRNA, respectively. One should note that these numbers were much smaller than those identified by Luding et al [3]. Thus, since these are feasible, our methodology has more power to get limited number of critical miRNAs/mRNAs.

### B. *Discrimination study between tumors and normal kidneys*

Since our methodology is unsupervised, one may wonder if we surely could get miRNA/mRNAs that are distinct between Wilms tumor and normal kidney. In order to confirm this point, we tried to discriminate Wilms tumor samples from normal kidney samples (Table I). It is obvious that discrimination is almost complete. For miRNA and mRNA, only one Wilms tumor sample was wrongly identified as normal kidney. Thus, we concluded that identified mRNA/miRNAs by PCA based unsupervised FE was surely distinct between normal kidneys and Wilms tumors.

**Table I.** Discrimination between normal kidneys and Wilms tumors. Row: prediction, column: true classes. The first PC loading for MRNA (*L* = 1) and the First Seven PC Loadings for miRNA (*L* = 7) were Employed for the Discrimination.

|  | mRNA |  | miRNA |  |
| --- | --- | --- | --- | --- |
|  | kidney | tumor | kidney | tumor |
| kidney | 4 | 1 | 4 | 1 |
| tumor | 0 | 27 | 0 | 61 |

### C. Identification of miRNA-mRNA pairs associated with reciprocal differential expression

In order to identify biologically meaningful miRNA-mRNA pairs, miRNA/mRNA significantly up-/down-regiulated between normal kidney and Wilms tumors were screened. *P*-values were attributed with *t* test to mRNAs and miRNAs, and were further adjusted by BH criterion. Then, those associated with adjusted P-values less than 0.01 were identified as significantly up-/down-regulated. Finally, among miRNA-mRNA pairs listed in mirTarBase [28], pairs of miRNA and mRNA associated with reciprocal differential expression are selected. Fig. 2 shows the miRNA-mRNA network composed of selected miRNA-mRNA pairs.

**Figure 2.**
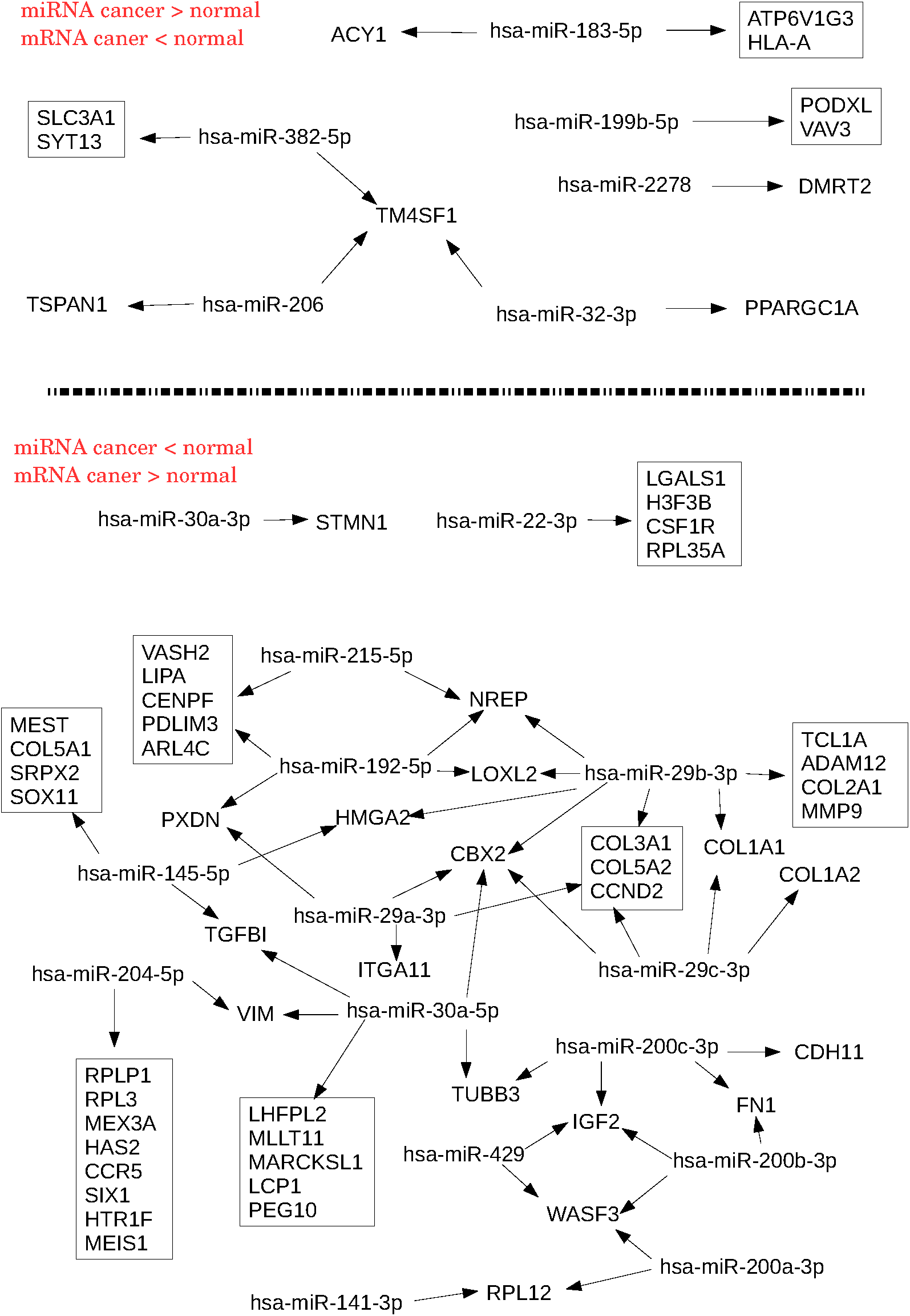
miRNA-mRNA network composed of identified miRNA-mRNA pairs.

It is obvious that the pairs of miRNA down-regulaged in tumor and mRNA up-regulated in tumor are more enhanced than those of miRNA up-regulated in tumor and mRNA down-regulated in tumor. The former has more miRNAs and its network are highly connected while the latter has less miRNAs and not connected network. It is reasonable, since Wilms tumor differentiated from kidney, gain of gene expression should be more probable than loss of that. If we compare them with Table 1 by Luding et al [28], feasibility of network in Fig. 2 is more enhanced. Luding et al listed the same number of 15 up-/down-regulated miRNAs in thier Table 1. Apparently, it is contradict to unbalanced identification of up-/down-regulated miRNAs in Fig. 2. However, more detailed inspection can reverse this impression; FC of miRNAs down-regulated in tumor is much larger than that for down-regulated miRNAs in Luding et al’s Table 1. The smallest FC in the former is as large as 7.01 while the largest FC among the latter is as small as 8.11 in their Table 1. This suggested that poor identification of miRNAs up-regulated in tumor in Fig. 2 is more reasonable than the first impression. In other words, our methodology correctly reflected the distinct importance between up-/down-regulated miRNAs in tumor samples. This suggested the usefulness of our methodology. In addition to this, it is remarkable that most of 15 miRNAs down-regulated in tumor listed in Luding et al’s Table 1 were identified in Fig. 2. Among 15 miRNAs, 10 miRNAs (miR-200a/b/c, 204, 141, 192, 429, 215, 30a, and 30a*) are in Fig. 2. Thus, our methodology correctly reproduced Luding et al’s subjective but biologically reasonable selections of miRNAs without tunable selection criteria.

## IV. DISCUSSION

### A. Significant negative correlation between miRNA-mRNA pairs

Although Luding et al [28] explicitly considered significant negative correlation between miRNA and mRNA, since our methodology was originally designed so as to be applied to unmatched data set [8], we did not require the significant negative correlations between miRNA-mRNA pairs explicitly. Although we required reciprocal differential expression, this did not always guarantee negative correlation, since samples are unbalanced between normal kidney and Wilms tumors (the number of Wilms tumor samples are much more than normal kidney); if there are no negative correlations within Wilms tumor samples, there may not be significant negative correlation between miRNA and mRNA. In order to confirm this point, we computed Pearson correlation coefficients between miRNA-mRNA pairs listed in Fig. 2. Then, mean values and attributed *P*-values computed by *t*-test; Null hypothesis to be rejected was that mean values of Pearson correlation coefficients are zero. Then, means and attributed *P*-values were −1.26 × 10^−1^ (P= 8.52 ×10^−3^) and −2.67 × 10^−1^ (*P* < 2.2 × 10^−16^, lower limit of numerical accuracy) for the pairs of up-regulated miRNAs and down-regulated mRNAs in tumor (upper half of Fig. 2) and those of down-regulated miRNAs and up-regulted mRNAs in tumor (lower half of Fig. 2), respectively. Thus, it is confirmed that mRNA and miRNAs expression are negatively correlated between miRNA-mRNA pairs listed in Fig.2.

### B. Biological feasibility of identified miRNA-mRNA pairs

Fig. 2 includes multiple mir-29s that extensively target collagen proteins which Wilms tumor massively synthesize [29] and whose inhibition was known to inhibit Wilms tumor [30], although miR-29s were missing in Luding et al’s Table 1 [28]. This suggests that the regulation of miR-29s are important potential therapy target and usefulness of our methodology. To our knowledge, since miR-29s were never therapy target of Wilms tumor, identification of this interaction may be important.

On the other hand, *IGF2* was targeted by three miRNAs, 200b/c and 429. Overexpression of *IGF2* was reported to be a disease cause [1]. Although dysregulation of miR-200s was reported to be associated with Wilms tumor [31], this is possibly the first report of miR-200s targeting *IGF2* in Wilms tumor, although miR-200s regulation of *IGF2* was reported in other ocasions [32].

miR-29s as well as miR-30 target *CBX2* which was once reported to be overexpressed in Wilms tumor with considering histone modification [33]. Thus, *CBX2* targeting miRNAs are potential new therapy target for Wilms tumor.

*NREP* was also targeted by three miRNAs, miR-219, 192 and 29b. Although the negative regulation of *NREP* by miR-29b was identified in cancer [34], *NREP* has never been regarded as therapy target of Wilms tumor.

Although *SIX1* was targeted by only one miRNA, miR-204, importance of its mutation in Wilms tumor was once reported [35].

Finally, *WASF3* targeted by three miRNAs (Fig. 2), together with *SIX1*, were once identified as two of 27 wilms tumor signature genes [36].

All of these above suggested the usefulness of our methodology to figure out the mechanism of Wilms tumor progression.

### C. Relationship with survival data

OncoLnc [37] can provide us the information if gene is significantly related to survival probabilities based on precomputed survival analyses. We have evaluated if genes targeted by multiple miRNAs in the lower half of Fig. 2 are significantly related to survival probabilities in various cancers (Table II). Interesting, excluding one exception *(NREp)*,all genes targeted by multiple miRNAs has relationship with survival probabilities in some cancer(s). More interestingly, most of them other than *WASF3* and *VIM* are related to survival probabilities of either of two renal cancers. Since OncoLoc unfortunately did not include Wilmes tumor, frequent relation to renal cancers’ survival probabilities also supported feasibility of our analyses. We have also noticed that lower grade glioma (LGG) is often listed in Table II. There are multiple studies [38], [39], [40] that report the enhanced expression of *WT1* in LGG. Since *WT1*, Wilms tumor 1, is expressive in Wilms tumor as its name says, frequent relationship between LGG and genes listed in Table II suggests that *WT1* is not only gene expressive in LGG as well as Wilms tumor. More studies are waited.

**Table II.**
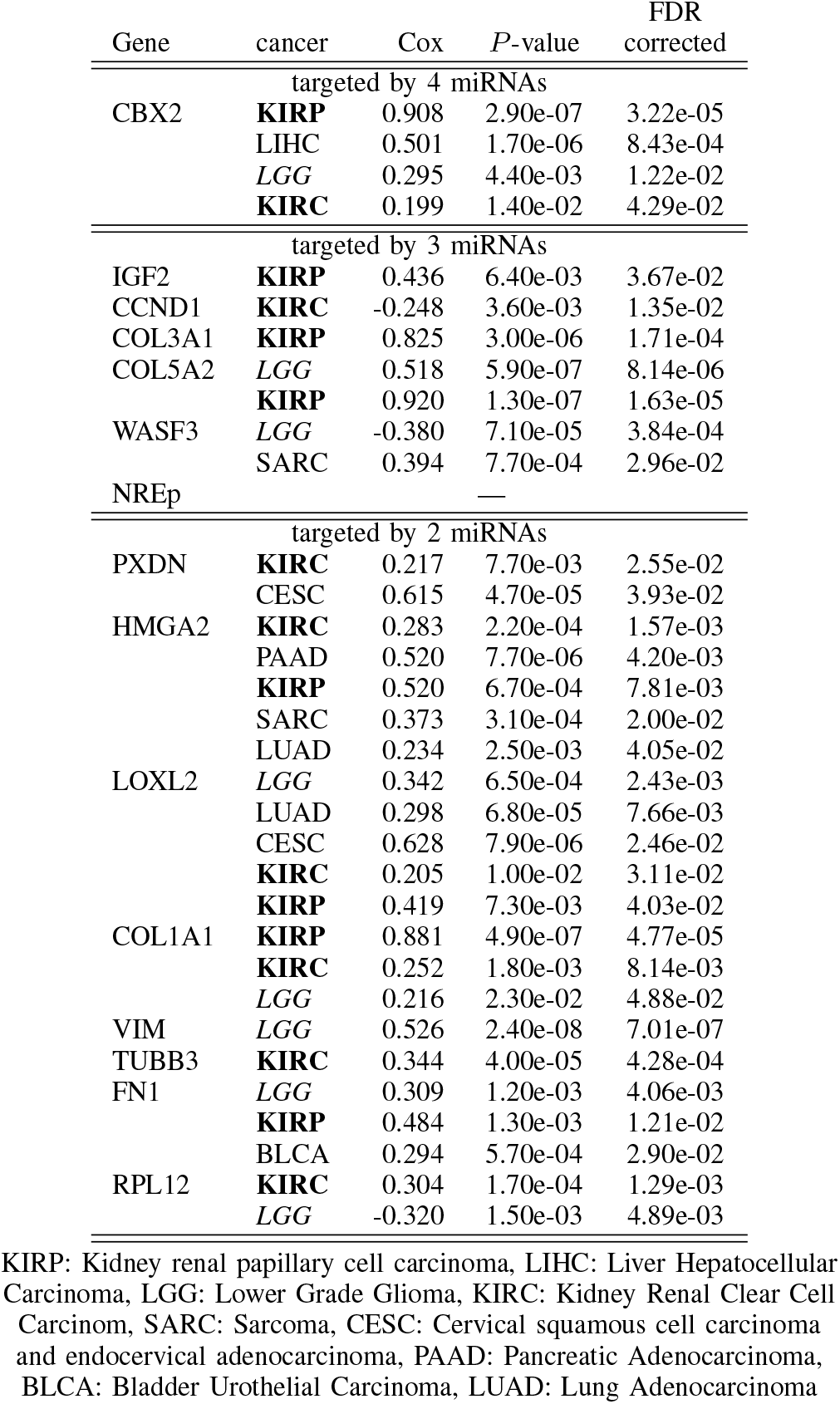
Significant Relationships to Survival Probabilities in Various Cancers Provided by Oncolnc. Those Associated with Corrected FDR < 0.05. Two Renal Cancers are in Bold. LGG was in Italic in Order to Emphasize the Association with Renal Cancers.

### D. Epigenetic landscape of Wilms tumor

Hohenstein et al [35] wrote in the recent review, “there are very few genes commonly mutated in Wilms tumor, and all show relatively low mutation frequencies.”. This is possibly another reason why people are interested in miRNAs; if mutation is not a potential cause, gene regulation which miRNAs mainly contribute to can be a primary factor. In this regard, we consider epigenetic landscape here. When uploading genes listed in lower half of Fig. 2, i.e., those up-regulated in tumor, to Enrichr [41] which lists many epigenetic features of genes, we can find many epigenetic feature enriched in these genes.

For example, SUZ12 and EHZ2 bindings to promoter region is enhanced in these genes (Adjusted P ranges from 1.418×^−10^ to 1.43 × 10^−5^ for SUZ12 and is 1.78 × 10^−6^ for EHZ2 in mouse embryonic stem cell (MESC) using CHEA2015, respectively. Adjusted *P* = 2.135 × 10^−4^ in SUZ12_CHEA using “ENCODE and ChEA Consensus TFs from ChIP-X”). SUZ12 and EZH2 are Ploycomb complex proteins and were reported to be recruited to suppress *Pax2* [42]. SUZ12 and EZH2 were also used for establishing H3K27me3 in Wilms tumor [43]. In actual, Enrichr reported H3K27me3 enhancement of these genes (Adjusted *P* = 2.02 × 10^−9^ in H3K27me3_kidney_mm9 using “ENCODE Histone Modifications 2015”).

Although Polycomb complex proteins BMI1 and EED were also reported to contribute to establish H3K27me3 in Wilms tumor [43], their enhanced binding to promoters was reported by Enrichr (Adjusted *P* = 2.13 × 10^−6^ in mouse neuronal progenitor cells (MNPC) for BMI1 Adjusted *P* = 2.25 × 10^−5^ in MESC for BMI1 using CHEA2015, respectively).

In addition to these, Hohenstein et al [35] discussed in detail Wilms tumor epigenetic anomaly of *IGF2* targeted by three miRNAs (Fig. 2).

Although there are no more reports about the relationship between enhanced TF bindings to promoter, in total as many as 83 TFs (Table III) were reported to significantly bind to promoter regions of genes up-regulated in tumor (Fig. 2). In the future, these enhanced TFs bindings to up-regulated genes in Wilms tumor may turn out to be a potential factor.

**Table III.** TFs reported in CHEA2015 to be enhanced (Adjusted *P*-values < 0.01) in genes up-regulated in tumor (Fig. 2)

|  |  |  |  |  |  |
| --- | --- | --- | --- | --- | --- |
| SUZ12 | EP300 | KLF4 | ZFP281 | OLIG2 | PHC1 |
| CCND1 | CEBPD | EZH2 | RNF2 | EED | TP53 |
| NRF2 | BMI1 | NFE2L2 | MTF2 | MEIS1 | TCF3 |
| AHR | SOX2 | JARID2 | WT1 | RUNX2 | RELA |
| POU3F2 | NR3C1 | NANOG | EKLF | ZNF217 | BRD4 |
| SETDB1 | SALL4 | FOXM1 | CTBP1 | SMARCA4 | ESR1 |
| SOX17 | ARNT | SMAD1 | TCF21 | CEBPB | YAP1 |
| PPARD | NACC1 | TOP2B | SCL | PAX3-FKHR | ATF3 |
| SMAD4 | MYC | ZFP42 | BACH1 | DROSHA | SRY |
| GFI1B | AR | NCOR1 | PPARG | EWS-FLI1 | JUND |
| NR0B1 | GATA2 | GATA1 | MNX1 | TCFAP2C | SOX9 |
| TAL1 | RAD21 | E2F1 | DMRT1 | POU5F1 | TP63 |
| KLF5 | FOXA2 | POU3F1 | RUNX1 | TCF4 | TET1 |
| E2F4 | EGR1 | SIN3B | HNF4A | FLI1 |  |

As described in the above, some enhanced TFs binding to promoter were related to histone modification. In actual, Enrichr reported extensive histone modification enrichment (Table IV) for genes up-regulated in tumor (Fig. 2). Aiden et al [33] reported that the amount of three histone modifications, H3K4me3, K3K36me3, and H3K27me3, were as much as in ESC. They concluded that this suggested that Wilms tumor keeps variability as much as ESC has.

These above suggested that miRNAs down-regulated in tumor possibly may target epigenetically important genes up-regulated in tumor. Since interaction between epigenetics and miRNAs were recently proposed [44], [45], this observation sounds feasible.

## V. CONCLUSION

In this paper, we applied the recently proposed PCA based unsupervised FE to published miRNA/mRNA expression profiles in Wilms tumor separately and identified limited number of miRNA-mRNA pairs. The pairs especially including mRNA up-regulated in tumor form highly connected network. mRNAs as well as miRNAs that form this highly connected network are often referred as factors related to Wilms tumor. In addition to this, mRNAs included in this network are highly enriched with TFs binding to promoter region as well as histone modifications. This suggested that identified down-regulated miRNAs possibly target mRNAs associated with epigenetic anomaly and its dysfunction contributed to Wilms tumor progression.

**Table IV.** Histone modification enrichment in “ENCODE Histone Modifications 2015” (Adjusted *P*-values < 0.01) in genes up-regulated in tumor (Fig. 2)

| tissue/cell line | $P$-values | adjusted $P$-values |
| --- | --- | --- |
| mouse (mm9) |  |  |
| H3K27me3 |  |  |
| small intestine | 2.49E-12 | 9.71E-10 |
| kidney | 1.08E-11 | 2.02E-09 |
| cerebellum | 1.55E-11 | 2.02E-09 |
| liver | 2.20E-10 | 2.15E-08 |
| testis | 9.82E-08 | 7.66E-06 |
| heart | 6.31E-07 | 3.51E-05 |
| thymus | 3.70E-06 | 1.44E-04 |
| ES-Bruce4 | 9.57E-05 | 1.97E-03 |
| megakaryocyte | 8.22E-05 | 1.97E-03 |
| spleen | 9.57E-05 | 1.97E-03 |
| erythroblast | 1.11E-04 | 2.16E-03 |
| G1E | 2.54E-04 | 4.71E-03 |
| H3K79me2 |  |  |
| myocyte | 1.98E-05 | 7.01E-04 |
| C2C12 | 1.65E-03 | 2.26E-02 |
| H3K36me3 |  |  |
| myocyte | 5.28E-05 | 1.72E-03 |
| C2C12 | 6.56E-05 | 1.97E-03 |
| H3K4me1 |  |  |
| brown adipose tissue | 9.57E-05 | 1.97E-03 |
| ES-Bruce4 | 4.19E-04 | 7.10E-03 |
| kidney | 1.65E-03 | 2.26E-02 |
| human (hg19) |  |  |
| H3K27me3 |  |  |
| mononuclear cell | 6.31E-07 | 3.51E-05 |
| endothelial cell of umbilical vein | 9.01E-05 | 1.97E-03 |
| CD14-positive monocyte | 1.39E-06 | 6.78E-05 |
| GM12878 | 2.48E-06 | 1.07E-04 |
| MCF-7 | 1.32E-03 | 2.14E-02 |
| H3K9me3 |  |  |
| CD14-positive monocyte | 9.57E-05 | 1.97E-03 |
| fibroblast of dermis | 4.19E-04 | 7.10E-03 |
| fibroblast of mammary gland | 1.68E-03 | 2.26E-02 |
| astrocyte | 1.65E-03 | 2.26E-02 |
| H4K20me1 |  |  |
| endothelial cell of umbilical vein | 1.65E-03 | 2.26E-02 |

## ACKNOWLEDGMENT

This study was supported by KAKENHI 26120528.

